# Constitutive upregulation of transcription factors underlies permissive bradyzoite differentiation in a natural isolate of *Toxoplasma gondii*

**DOI:** 10.1101/2024.02.28.582596

**Authors:** Jing Xia, Yong Fu, Wanyi Huang, L. David Sibley

## Abstract

*Toxoplasma gondii* bradyzoites play a critical role in pathology due to their long-term persistence in intermediate hosts and their potential to reactivate, resulting in severe diseases in immunocompromised individuals. Currently there is no effective treatment for eliminating bradyzoites. Hence, better *in vitro* models of *T. gondii* cyst development would facilitate identification of therapeutic targets for bradyzoites. Herein we characterized a natural isolate of *T. gondii*, called Tg68, which showed slower *in vitro* replication of tachyzoites, and permissive bradyzoite development under stress conditions *in vitro*. Transcriptional analysis revealed constitutive expression in Tg68 tachyzoites of the key regulators of bradyzoite development including *BFD1*, *BFD2*, and several AP2 factors. Consistent with this finding, Tg68 tachyzoites expressed high levels of bradyzoite-specific genes including *BAG1*, *ENO1*, and *LDH2*. Moreover, after stress induced differentiation, Tg68 bradyzoites exhibited gene expression profiles of mature bradyzoites, even at early time points. These data suggest that Tg68 tachyzoites exist in a pre-bradyzoite stage primed to readily develop into mature bradyzoites under stress conditions *in vitro*. Tg68 presents a novel model for differentiation *in vitro* that will serve as a useful tool for investigation of bradyzoite biology and development of therapeutics.

**Significance:** *Toxoplasma gondii* is a widespread protozoan that chronically infects ∼30% of the world’s population. *T. gondii* can differentiate between the fast-growing life stage that causes acute infection and the slow-growing stage that persists in the host for extended periods of time. The slow-growing stage cannot be eliminated by the host immune response or currently known antiparasitic drugs. Studies on the slow-growing stage have been limited due to the limitations of *in vivo* experiments and the challenges of *in vitro* manipulation. Here, we characterize a natural isolate of *T. gondii*, which constitutively expresses factors that drive development and that is permissive to convert to the slow-growing stage under stress conditions *in vitro*. The strain presents a novel *in vitro* model for studying the chronic phase of toxoplasmosis and identifying new therapeutic treatments for chronic infections.

## Introduction

*Toxoplasma gondii* is a widespread opportunistic protozoan parasite capable of infecting a broad range of warm-blooded hosts, including wild, companion, and domesticated animals. It is a frequent cause of zoonotic infection in human and it is estimated that approximately a third of the world’s population is chronically infected with *T. gondii*(1, 2). Although most cases of infection are associated with mild symptoms in immunocompetent individuals, toxoplasmosis can be life-threatening in immunocompromised individuals(3, 4). For example, *T. gondii* is reported to cause encephalitis, chorioretinitis, and pneumonitis in immunocompromised patients(5–7). Maternal infections with *T. gondii* during pregnancy can also lead to transplacental transmission, causing abortion, fetal death, and cerebral and/or ocular abnormalities in infected newborns or infants(8, 9).

*T. gondii* undergoes a two-host life cycle(10). Its definitive host is the Felidae (cat) family, where the parasite replicates sexually in the intestine to produce oocysts, which are excreted in feces and become infective after sporulation(10). When ingested by an intermediate host, which includes a wide range of warm-blooded animals, sporozoites contained in the oocysts are liberated and invade the intestinal epithelium, where they differentiate into tachyzoites(3). Tachyzoites spread rapidly throughout the host’s body and result in inflammatory response and tissue destruction, thus causing clinical manifestations of diseases(1, 3). In response to stress and immune pressure, tachyzoites differentiate into slow-growing bradyzoites, which persist within cysts for extended periods of time, typically in brain and muscle(11). Once considered dormant, more recent studies have shown that bradyzoites within tissue cysts replicate periodically and asynchronously to support the expansion of cyst size *in vivo*(12). Cysts eventually rupture, releasing bradyzoites that are either eliminated by the immune system or that give rise to daughter cysts(13). Patients receiving immunosuppressive therapy and congenitally infected individuals are at risk of reactivation of pre-existing latent infection(14). During reactivation, the conversion of bradyzoites into tachyzoites results in dissemination of the parasite that can lead to severe pathology in immunocompromised hosts(11). Reactivation of chronic *T. gondii* infection is one of the most common causes of central nervous system complications in immunocompromised hosts(5). Therefore, the long-term persistence of bradyzoite-containing cysts in hosts and the potential reactivation of chronic infections play a key role in *T. gondii* pathogenesis. However, currently accepted treatments for *T. gondii* infection only target the acute stage of the parasite, and there is no effective clinically approved treatment for eradicating *T. gondii* bradyzoites within tissue cysts(15).

Studies on the development of bradyzoites within tissue cysts in vivo have been hampered by the difficulty in obtaining large quantities of tissue cysts from *in vivo* models. One recent development that addresses this concern in part is the characterization of a derivative of ME49 that is strictly passed *in vivo* and which produces large numbers of tissue cysts(16). However, this phenotype is compromised by even short-term passage *in vitro*, rendering the strain less suitable for genetic manipulation. In contrast, *in vitro* culture systems, where *T. gondii* bradyzoites can be induced by a variety of treatments including alkaline stress(17), allow for consistent and scalable productions of cysts. However, previous studies have pointed out that such *in vitro* differentiated bradyzoites may not express the full range of features, nor the full repertoire of gene expression changes, seen in mature cysts *in vivo*(18–20). Typically, the cyst-forming type II strain ME49 has been used *in vitro* for bradyzoite research; however, the cysts produced by this strain *in vitro* always contain bradyzoites that are only partially differentiated(21–23). The hybrid nature of bradyzoites and tachyzoites in culture can result in lysis of host cell monolayer by the rapidly replicating tachyzoites, thereby disrupting the culture system needed for long-term maintenance of cysts. In addition, the different characteristics and behaviors between bradyzoites and tachyzoites introduce variations in gene expression patterns and responses to treatments, which cannot be distinguished without transcriptomic or proteomic analyses at single-cell resolution.

Here, we described a new isolate of *T. gondii* called Tg68, which shows constitutive upregulation of bradyzoite developmental regulators in tachyzoites and is permissive for mature bradyzoite differentiation under stress conditions *in vitro*. As such, Tg68 may serve as a useful tool for investigating the mechanisms of *T. gondii* stage conversion *in vitro* and identifying potential therapeutic agents that target bradyzoites.

## Results

### Tg68 parasites exhibit distinct growth phenotypes *in vitro* compared to ME49 parasites

Tg68 was initially isolated from a patient with congenital toxoplasmosis by Jack Frenkel and was identified as a Type II strain by polymerase chain reaction-restriction fragment length polymorphisms (PCR-RFLP)(24). Tachyzoites of the isolate were preserved by storage in liquid nitrogen and subsequently recovered in a human foreskin fibroblasts (HFFs) culture following thawing. Although Tg68 tachyzoites exhibited a typical lytic pattern of growth on HFF monolayers, we noticed that Tg68 tachyzoites replicated slower, resulting in a longer incubation time between passages compared to Type II strain ME49 tachyzoites. To assess the *in vitro* properties growth of Tg68 tachyzoites, we compared the size of parasitophorous vacuoles (PVs) of Tg68 tachyzoites with that of ME49 tachyzoites at 24, 48, and 72 hr post inoculation (Fig. 1A). Based on staining of the surface antigen SAG1, Tg68 tachyzoites had smaller PVs compared to ME49 tachyzoites (Fig. 1A). Quantitative image analysis revealed that the average PV diameter of Tg68 tachyzoites was significantly smaller than that of ME49 tachyzoites at all three time points (Fig. 1B). When cultured in HFFs under alkaline stress, which induces *T. gondii* bradyzoite differentiation *in vitro*(17), Tg68 robustly formed BAG1-positive bradyzoites-containing cysts that were stained with *Dolichos biflorus* Agglutinin (DBA) lectin that recognizes α-1,3-N-acetylgalactosamine linkages in the cell wall (Fig. 1C and 1D). In contrast, at the relatively high MOI used here, ME49 cultures showed a significantly lower yield of cysts and contained many SAG1-positive tachyzoites under the identical alkaline stress condition (Fig. 1C and 1D). Together, these data show that Tg68 tachyzoites exhibit slow replication *in vitro*, and a markedly high rate of bradyzoite differentiation under alkaline pH conditions. Remarkably, this trait was also stable at high MOI, leading to much higher yields of bradyzoites from *in vitro* differentiated cultures.

**Figure 1.**
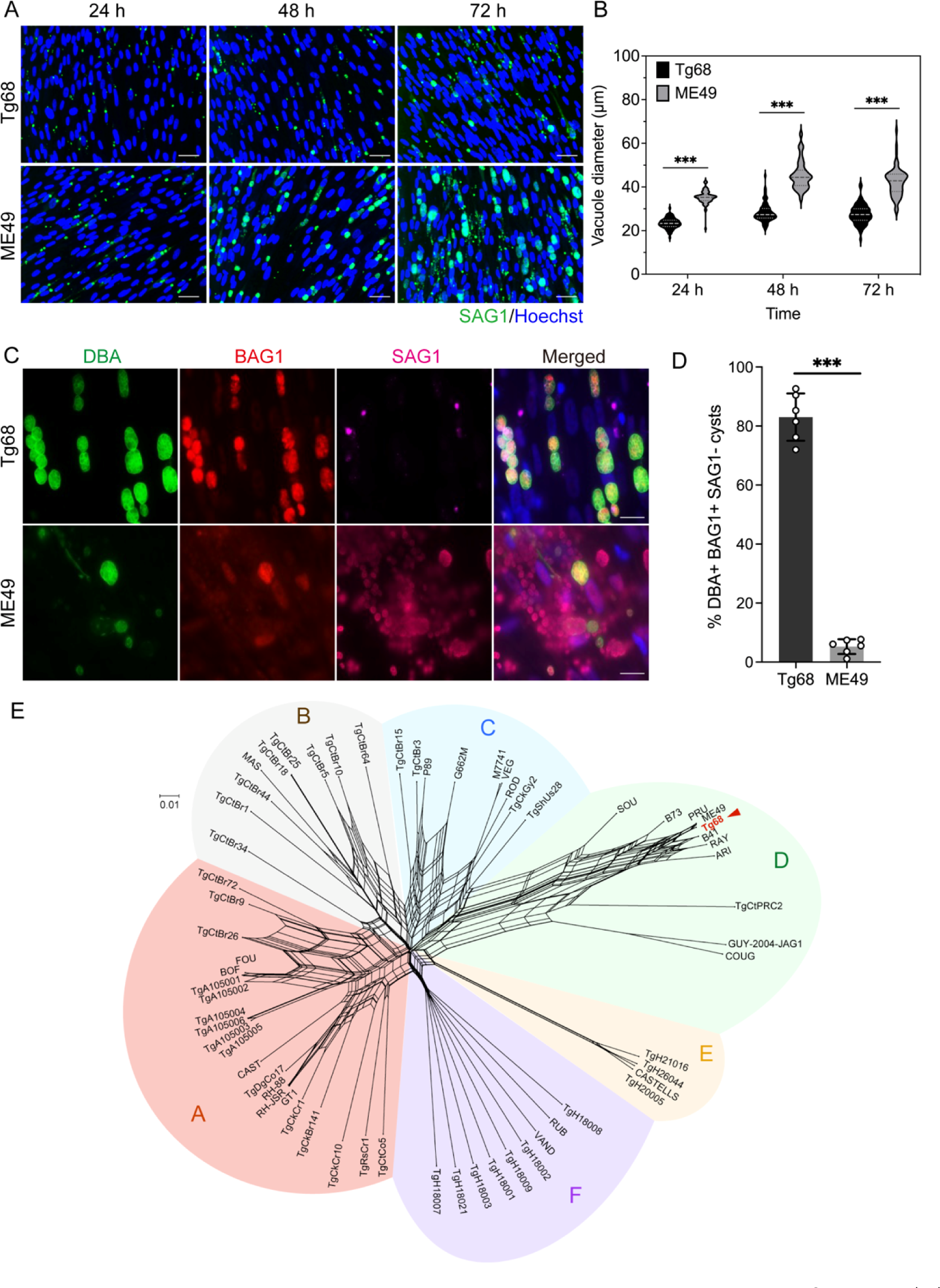
*In vitro* phenotypes and genetic characterization of Tg68. (A) HFFs were infected with Tg68 or ME49 tachyzoites at an MOI of 5 for 24, 48, or 72 hr. Cells were fixed and IF staining was performed with anti-SAG1 rabbit antibody and goat anti-rabbit secondary antibody conjugated to Alexa Fluor 647. Hoechst 33342 was used to stain DNA. Vacuoles were visualized using a Cytation 3 Imager and representative images are shown. Scale bar, 50 μm. (B) Violin plot representing diameters of vacuoles of Tg68 or ME49 tachyzoites at indicated timepoints. Data are combined from three independent experiments in which at least 5,000 vacuoles per strain were measured in each experiment. In violin plot, the middle line indicates median, the upper line indicates the 75^th^ percentile, and the lower line indicates the 25^th^ percentile. ***, Student’s t-test, *P* < 0.001. (C) HFFs infected with Tg68 or ME49 tachyzoites were cultured under alkaline pH condition (pH = 8.2, ambient CO_2_) for 7 days. Cells were fixed and stained with anti-BAG1 mouse mAb 8.25.8 and anti-SAG1 rabbit antibody, followed by goat anti-mouse secondary antibody conjugated to Alexa Fluor 568 and goat anti-rabbit secondary antibody conjugated to Alexa Fluor 647. Dolichos biflorus agglutinin (DBA) conjugated to Alexa Fluor 488 was used to stain the cyst wall and Hoechst 33342 was used to stain DNA. Scale bar, 20 μm. (D) Percentage of DBA-positive BAG1-positive SAG1-negative cysts of Tg68 or ME49. Mean ± SE was plotted for 6 independent experiments, in which at least 100 vacuoles/cysts were counted in each experiment. ***, Student’s t-test, *P* < 0.001. (E) Neighbor-net analysis of Tg68 and 62 *T. gondii* isolates based on 804,624 SNPs. Major clades of *T. gondii* are indicated in different colors. Scale bar, number of SNPs per site.

### Genetic characterization of Tg68

Although Tg68 is known to be a type II strain based on RFLP analysis, to attain a more comprehensive understanding of its genomic composition, we conducted whole genome sequencing using Illumina short-read Next Generation Sequencing. We compared single nucleotide polymorphisms (SNPs) identified in Tg68 to 62 previously charactered genomes that were part of a comparative genome analyses of diverse *T. gondii* lineages(25). Neighbor-net analysis of Tg68 and 62 *T. gondii* isolates based on 804,624 single nucleotide polymorphisms (SNPs) at shared positions revealed that Tg68 is closely related to strains contained in clade D, including the commonly used lab strain ME49, as well as closely related PRU, B41, and RAY strains (Fig. 1E). A total of 4,918 SNPs, 227 insertions, and 216 deletions were detected in Tg68 genome when compared to ME49 genome. Of these, 16 variants were predicted to have a high functional impact, including SNPs that caused start loss, acceptor splice site variant, frameshift, stop gain, and stop loss (Table S1). Among the high-impact variants, the initiating ATG was lost in guanylyl cyclase (*TgGC*), which could potentially affect the PKG mediated invasion and egress pathways(26). *TgGC* is an essential gene, suggesting an alternative downstream ATG may be used instead (Fig. S1), although we have not interrogated this experimentally. We also noted a frameshift variant in elongation factor thermo stable (*EF-Ts*) that could have a potential impact on mitochondrial translation(27) (Fig. S1). The majority of other changes were in surface secretory or hypothetical unknown proteins (Table S1). Although these alterations might affect various aspects of the biology, our analysis did not identify any variants of genes known to be directly associated with bradyzoite development.

### Transcriptional analysis of Tg68 tachyzoites and bradyzoites

To further characterize bradyzoite differentiation in Tg68, we examined the transcriptional changes throughout the differentiation process by RNA sequencing (RNAseq). To monitor stage conversion of Tg68 *in vitro*, we developed a dual reporter strain of Tg68 that expressed fluorescent tags in a stage-specific manner, where the SAG1 promoter drives GFP expression and the BAG1 promoter drives mCherry expression (Fig. 2A). The expression of fluorescent tags was tested by monitoring expression by immunofluorescence (IF) staining at different times after induction by culture at alkaline pH (Fig. 2B). Under normal culture conditions (non-alkaline), vacuoles were negative for DBA and GFP positive, reflecting expression from the SAG1 promoter. At days 7, 14, and 21 post induction, DBA-positive cysts containing mCherry-positive parasites were robustly detected (Fig. 2B). Quantification of fluorescence intensities of GFP and mCherry revealed high GFP expression in DBA-negative vacuoles under normal culture conditions where tachyzoites predominate, vs. high mCherry expressions in DBA-positive cysts under alkaline stress conditions (Fig. 2C). We also sought a different method to induce differentiation based on previous work showing that high glucose inhibits bradyzoite development in HFF cells(28), and the finding that *T. gondii* can grow on glutamine as an alternative energy source(29). We also observed strong induction of bradyzoites when the reporter parasites were cultured in glucose-free conditions containing high levels of glutamine, which induced tachyzoite-to-bradyzoite conversion (Fig. 2C). Although the extent of differentiation appeared similar at 14 and 21 days post induction (dpi) to that of 7 dpi (Figs. 2B and 2C), we were interested in interrogating other genes in addition to the two promoters used. Consequently, we sequenced mRNAs from Tg68 after growth under normal conditions (tachyzoites) and after alkaline induction for 7 vs. 21 days.

**Figure 2.**
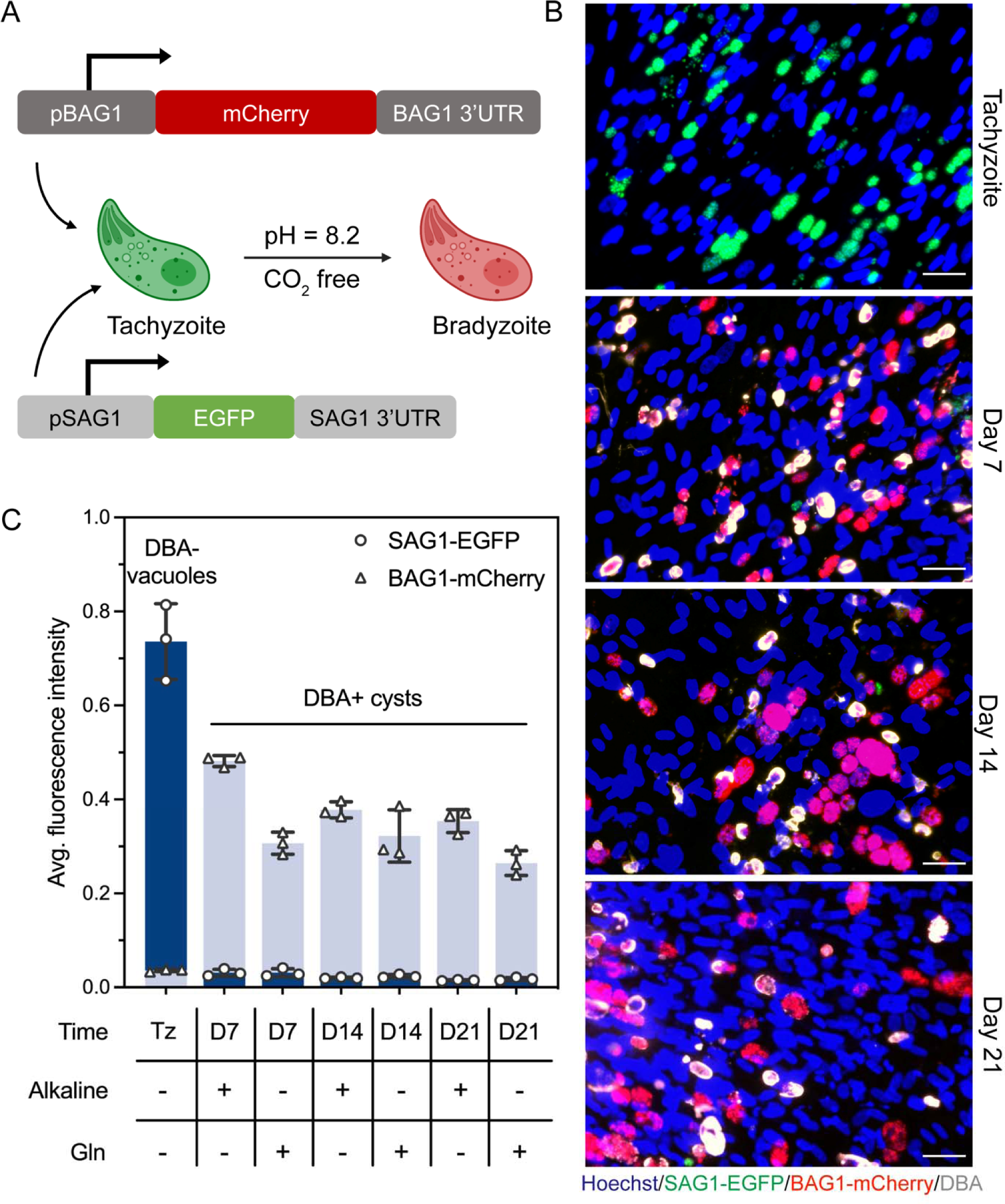
Tg68 bradyzoite development *in vitro*. (A) Generation of a stage specific dual fluorescent reporter strain of Tg68. The SAG1 promoter (pSAG1) drives EGFP expression and the BAG1 promoter (pBAG1) drives mCherry expression. (B) HFFs infected with SAG1-EGFP BAG1-mCherry Tg68 tachyzoites were cultured under alkaline pH conditions (pH = 8.2, ambient CO_2_) for 7, 14, or 21 days. Cells were fixed and DBA conjugated to Alexa Fluor 488 was used to stain the cyst wall. DNA was stained with Hoechst. Scale bar, 50 μm. (C) HFFs infected with SAG1-EGFP BAG1-mCherry Tg68 tachyzoites were cultured under alkaline pH conditions (pH = 8.2, ambient CO_2_) or glucose-free conditions (glucose free, 10 mM glutamine, pH = 7.2, 5% CO_2_) for 7, 14, or 21 days. Cells were fixed and DBA conjugated to Alexa Fluor 488 was used to stain the cyst wall. Average fluorescence intensities of EGFP and mCherry in tachyzoites (Tz) in DBA-negative vacuoles and in parasites in DBA+ cysts were determined using CellProfiler. Mean ± SE plotted for 3 independent experiments in which at least 4,500 vacuoles and 1,000 cysts were quantified in each experiment.

To efficiently capture bradyzoite stages and eliminate any residual tachyzoites from the differentiated cultures, we adopted a previously described magnetic bead-based protocol(30) to purify Tg68 cysts and prepare bradyzoite samples for RNAseq (Fig. 3A and Methods). Following the initial isolation of *in vitro* cysts from host cells, microscopic examination revealed the presence of cysts and tachyzoites (Fig. 3A). The application of anti-SAG1 antibody and protein G magnetic beads effectively removed tachyzoites, and cysts were subsequently enriched using biotinylated DBA and streptavidin magnetic beads (Fig. 3A). The purified cysts were then extracted and prepared for RNAseq. Transcriptional analysis revealed significant alterations in gene expression profiles in comparing day 7 bradyzoites vs. tachyzoites and day 21 bradyzoites vs. tachyzoites (Fig. 3B, left panel). We identified 1,560 upregulated genes and 812 downregulated genes in day 7 bradyzoites when compared to tachyzoites (FDR-adjusted *P* value ≤ 0.05 and absolute fold change ≥ 2) and 1,305 upregulated genes and 848 downregulated genes in the comparison day 21 bradyzoites vs tachyzoites (FDR-adjusted *P* value ≤ 0.05 and absolute fold change ≥ 2) (DataSet 1). The tachyzoite-specific gene *SAG1* was significantly downregulated in day 7 and day 21 bradyzoites (Fig. 3B and DataSet 1). The canonical bradyzoite-specific genes *BAG1*, *ENO1*, *LDH2*, and *SRS9* were among the top 20 differentially expressed genes (DEGs) with the greatest Euclidean distances in both day 7 and day 21 bradyzoites (Fig. 3B, right panel). To identify the time-dependent difference in gene expression profiles between the early stage bradyzoites and the late stage bradyzoites, we compared the gene expressions in Tg68 tachyzoites and day 7 bradyzoites to day 21 bradyzoites in the present study, and the comparison revealed a subset of genes only showed a significant upregulation in the late stage bradyzoites (Fig. 3C). Interestingly, among these genes, 12 were ribosomal proteins (Fig. 3C), which initially seemed at odds with the idea that bradyzoites are semi-dormant. However, the enrichment of ribosomal proteins is consistent with a previous study that used isobaric tags for relative and absolute quantification (iTRAQ)-based quantitative proteomics analysis of *T. gondii* life stages showing that 46 out of the 79 *T. gondii* ribosomal proteins were upregulated in day 28 *in vivo* bradyzoites compared to tachyzoites(31). These findings indicate that bradyzoites are metabolically active, and not dormant stages. To assess the maturity of Tg68 *in vitro* bradyzoites, we obtained the ME49 day 28, day 90, and day 120 *in vivo* bradyzoites transcriptional data from a previous study(18) and compared it to the Tg68 *in vitro* bradyzoites transcriptional data. The expression levels of *BAG1*, *ENO1*, and the cyst wall protein *CST1* in both Tg68 day 7 and day 21 *in vitro* bradyzoites were comparably high to those in ME49 day 28, day 90, and day 120 *in vivo* bradyzoites(18), suggesting that Tg68 *in vitro* bradyzoites exhibit similar maturity to that of the late stage *in vivo* bradyzoites (Fig. S2). While the expression levels of *SRS9* and *LDH2* in Tg68 *in vitro* bradyzoites were slightly lower than those in ME49 *in vivo* bradyzoites from Garfoot *et al*., 2019(18), they were remarkedly elevated compared to tachyzoites (Fig. S2).

**Figure 3.**
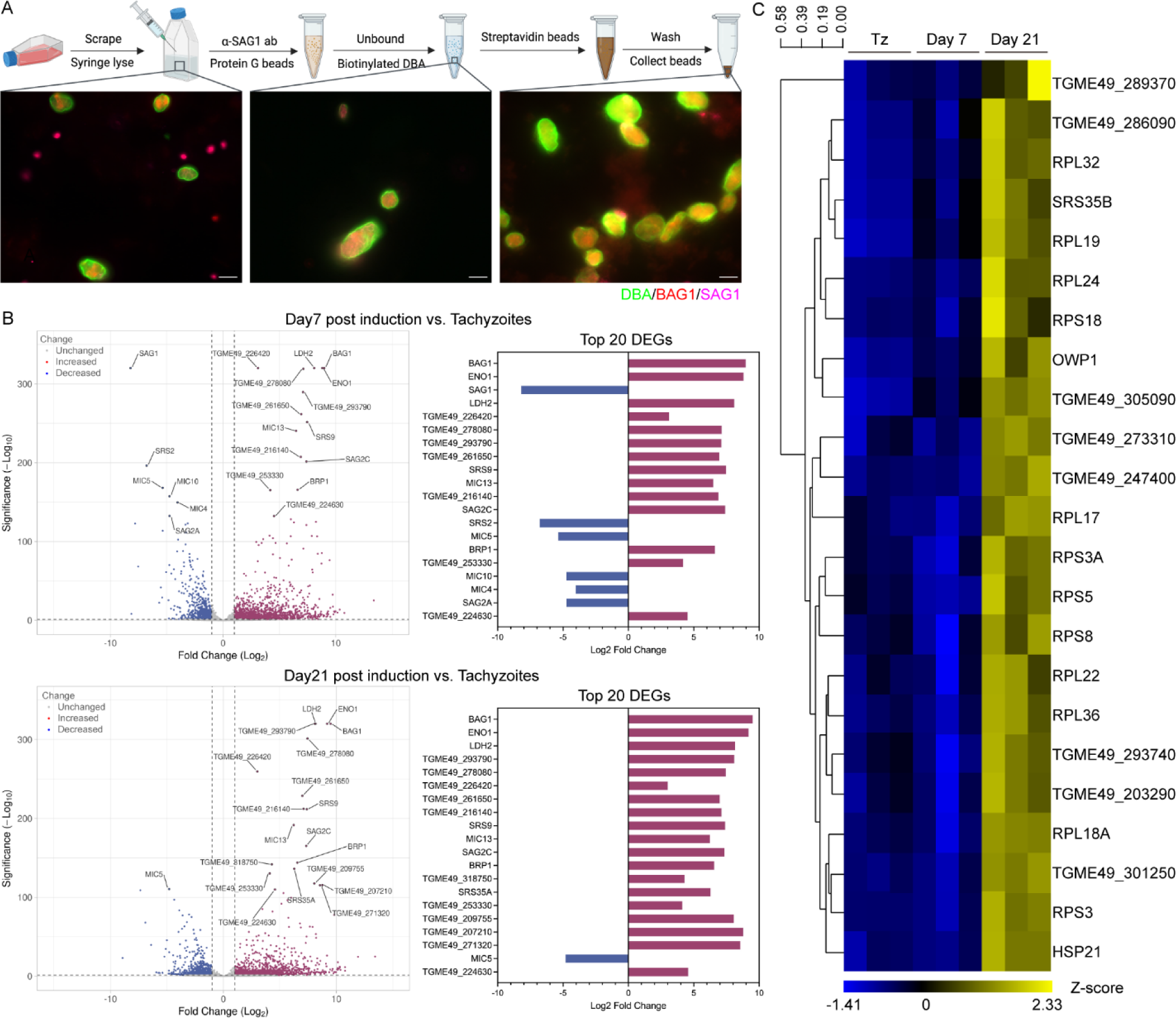
Transcriptional analysis of Tg68 tachyzoites and bradyzoites. (A) Schematic diagram of purification of Tg68 cysts induced *in vitro*, as described in the “*In vitro* cyst purification” section. Representative images of the samples after syringe lysis (left), tachyzoite removal (middle), and the enriched purified cysts (right) are shown. The cyst wall was stained with DBA conjugated to Alexa Fluor 488, BAG1 with anti-BAG1 mouse mAb 8.25.8 and goat anti-mouse secondary antibody conjugated to Alexa Fluor 568, and SAG1 with anti-SAG1 rabbit antibody and goat anti-rabbit secondary antibody conjugated to Alexa Fluor 647. Scale bar, 10 μm. (B) Volcano plot of DEGs (log_2_ fold-change > 1 or log_2_ fold-change < -1, and FDR adjusted *P* value ≤ 0.05) between Tg68 bradyzoites induced at pH 8.2 for 7 or 21 days and Tg68 tachyzoites (left panel). Fold change of the top 20 DEGs with the greatest Euclidean distances (right panel). (C) Heatmap representing the transcriptional abundance of mature bradyzoite markers during Tg68 bradyzoite development. Tz, tachyzoite.

### Bradyzoite development regulators are transcriptionally altered in Tg68 tachyzoites

The distinct phenotypes observed in Tg68 when compared to ME49 prompted us to investigate the transcriptional differences between Tg68 and ME49 tachyzoites and bradyzoites. Comparison of Tg68 and ME49 tachyzoites revealed a lower mRNA level of *SAG1* and a higher mRNA level of *BAG1* in Tg68 tachyzoites (Fig. 4A). Apetala2 (AP2) transcription factors are known to play important roles in regulating stage conversion in Apicomplexa(32), and six AP2 factors have been associated with bradyzoite development in *T. gondii*(33). Of these, AP2I-V4, which is known to suppress *T. gondii* bradyzoite development(34), exhibited a decreased transcription in Tg68 tachyzoites when compared to ME49 tachyzoites (Fig. 4A). Additionally, AP2XI-I2 has been shown to work with the microrchidia (MORC) transcriptional repressor complex and another AP2 transcription factor AP2IX-4, repressing bradyzoite differentiation *in vitro*(35). Our analysis showed that both *AP2XI-I2* and *AP2IX-4* were transcriptionally downregulated in Tg68 tachyzoites with comparison to ME49 tachyzoites (Fig. 4A). As well, AP2XI-4 is an activator of bradyzoite development(36), and the mRNA level of *AP2XI-4* was higher in Tg68 tachyzoites than that in ME49 tachyzoites (Fig. 4A). Finally, AP2IX-9 has been identified as a repressor of bradyzoite formation and it is highly expressed during early bradyzoite development(37). An increase in *AP2IX-9* transcription was detected in Tg68 tachyzoites compared to ME49 tachyzoites (Fig. 4A).

**Figure 4.**
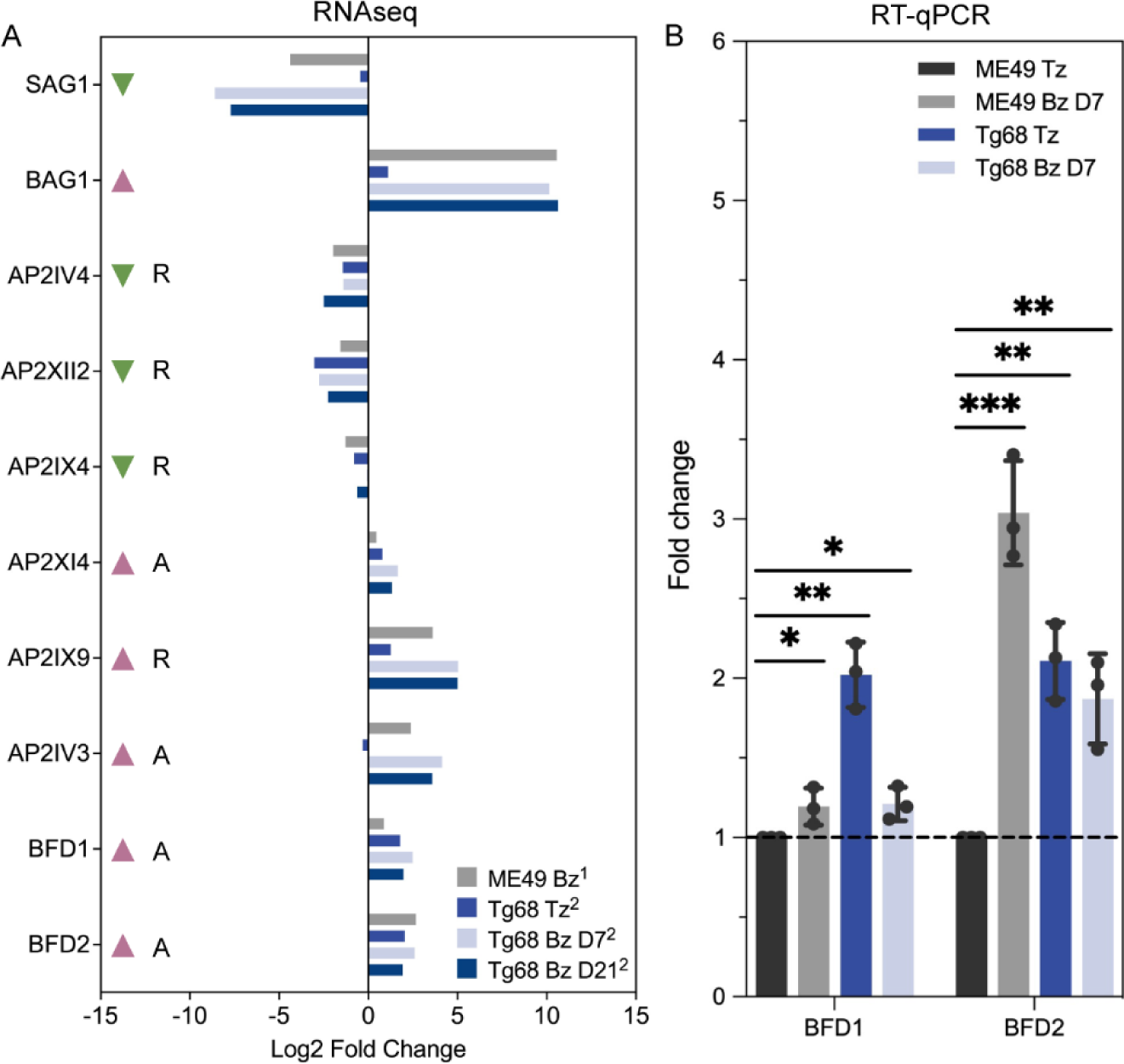
Expressions of bradyzoite development regulators in Tg68 tachyzoites and bradyzoites. (A) Fold change of known bradyzoite development-related genes in ME49 bradyzoites (ME49 Bz), Tg68 tachyzoites (Tg68 Tz), Tg68 Day 7 bradyzoites (Tg68 Bz D7), and Tg68 Day 21 bradyzoites (Tg68 Bz D21), compared to ME49 tachyzoites. Red and green arrows denote genes found to be over-expressed or under-expressed during bradyzoite development based on previous studies. A, activator of bradyzoite gene expression. R, repressor of bradyzoite gene expression. ^1^Data was obtained from Waldman *et al*., 2020(38). ^2^Data was from this study. (B) *BFD1* and *BFD2* mRNA levels were assessed by RT-qPCR, normalized to *GCN5B*. *, Student’s t-test, *P* < 0.05. **, Student’s t-test, *P* < 0.01. ***, Student’s t-test, *P* < 0.001.

Consistent with the observed upregulation of AP2 activators and downregulation of AP2 repressors in Tg68 tachyzoites, the master transcription regulator of bradyzoite development called Bradyzoite Deficient 1 (BFD1)(38) and its translation regulator Bradyzoite Deficient 2 (BFD2)(39)/Regulator of Cystogenesis 1 (ROCY1)(40) showed increased expression in Tg68 tachyzoites when compared to ME49 tachyzoites (Fig. 4A). Reverse transcription-quantitative polymerase chain reaction (RT-qPCR) results revealed consistent patterns of differential expression for *BFD1* and *BFD2* between ME49 and Tg68 tachyzoites (Fig. 4B). In Tg68 tachyzoites, *BFD1* mRNA exhibited a significant upregulation with an average fold change of 2 compared to ME49 tachyzoites (Fig. 4B). Similarly, *BFD2* transcription presented a 2.1-fold upregulation in Tg68 tachyzoites compared to ME49 tachyzoites (Fig. 4B). Taken together, our analysis revealed significant differences in the expression profiles of the key regulators of *T. gondii* bradyzoite development between Tg68 tachyzoites and ME49 tachyzoites, which is in line with the observed phenotypic differences between them.

### BFD1 and BFD1-regulated genes are upregulated in Tg68 tachyzoites

In the presence of alkaline stress, BFD1 binds to the promoters of stage-specific genes and induces a transcriptional program for bradyzoite differentiation(38). As the data above shows, BFD1 transcript levels are intrinsically higher in Tg68 tachyzoites (Fig. 4), and therefore we hypothesized that the transcription of BFD1-regulated genes would also be upregulated in Tg68 tachyzoites. To test this hypothesis, we compared the fold changes of BFD1-regulated genes in BFD1 overexpression ME49 parasites based on previous studies(38), to the corresponding levels in Tg68 tachyzoites (Fig. 5A). Seventy-five out of the 98 previously shown to be BFD1-upregulated genes (data points with Y values > 0) showed increased transcriptions in Tg68 tachyzoites compared to ME49 tachyzoites (Fig. 5A, red data points). Included in this set of upregulated genes are the bradyzoite-specific genes *BAG1*, *SRS9*, *ENO1*, and *CST1*, and the bradyzoite development regulators *BFD2* and *AP2IX-9* (Fig. 5A, red data points). Conversely, 14 previously shown to be BFD1-downregulated genes (data points with Y values < 0) exhibited reduced transcriptional levels in Tg68 tachyzoites with comparison to ME49 tachyzoites (Fig. 5A, blue data points).

**Figure 5.**
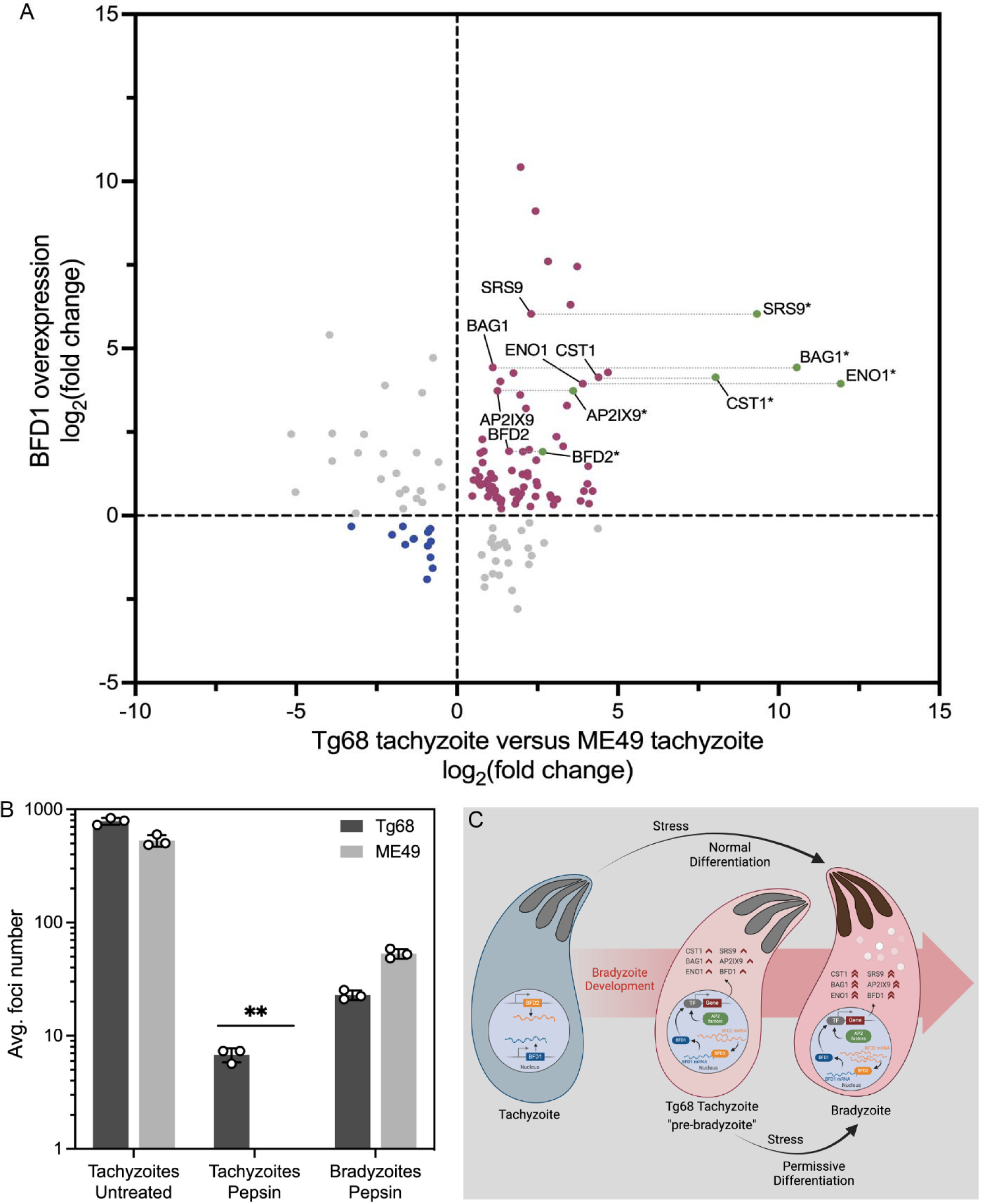
BFD1 and BFD1-regulated genes are upregulated in Tg68 tachyzoites. (A) Comparison of BFD1-regulated genes in BFD1 overexpression parasites (Waldman *et al*., 2020(38)) and Tg68 tachyzoites (present data) (significantly regulated genes with FDR adjusted *P* values of ≤ 0.05 in Tg68 tachyzoites compared to ME49 tachyzoites). *, fold changes correspond to ME49 bradyzoites versus ME49 tachyzoites. The ME49 transcriptional data used to generate this figure was obtained from Waldman *et al*., 2020(38). (B) HFFs infected with 1 x 10^4^ untreated tachyzoites, acid pepsin treated tachyzoites, and acid pepsin treated *in vitro* bradyzoites were cultured for 5 days. The number of foci were counted from 3 wells of a 24-well plate per sample and the average number of foci was determined. Mean ± SE was plotted for 3 independent experiments. **, Student’s t-test, *P* < 0.01. (C) Schematic model representing the “pre-bradyzoite” state of Tg68 tachyzoite. Constitutive upregulation of BFD1, BFD2, AP2 factors, and additional bradyzoite-specific genes in Tg68 tachyzoite makes the strain permissive for bradyzoite development and cyst formation under stress conditions *in vitro*. The schematic model was generated using BioRender. Gene set details are provided in Methods. See also DataSet 2.

The above data are consistent with induction of *BFD2* in Tg68 by a factor of 4.1- and 2.1-fold as shown by RNAseq and RT-qPCR, respectively (Fig. 4). In contrast, the expression of BFD2 showed higher levels in ME49 bradyzoites, with fold changes of 6.3 and 3.0 in RNAseq and RT-qPCR, respectively (Fig. 4). This partial upregulation of *BFD2* transcription in Tg68 tachyzoites was consistent with the observation that the fold changes of the BFD1-upregulated genes in ME49 bradyzoites were higher than that in Tg68 tachyzoites (Fig. 5A, green vs red data points).

In contrast to tachyzoites, bradyzoites exhibit resistance to low pH gastric digestion, allowing for oral infectivity in definitive and intermediate hosts(41). The resistance to acid pepsin has been used as a criterion to differentiate between tachyzoites and bradyzoites(41), and to evaluate the maturity of *in vitro* bradyzoites(42, 43). To determine whether Tg68 tachyzoites also exhibit resistance to acid pepsin, we treated the tachyzoites of Tg68 and ME49 strains with acid pepsin for 10 min and compared their viabilities by focus forming assay, along with the untreated tachyzoites and acid pepsin treated *in vitro* bradyzoites as controls. The untreated Tg68 and ME49 tachyzoites grew rapidly by forming more than 400 foci at day 5 post infection (Fig. 5B). The average foci number of acid pepsin treated Tg68 and ME49 bradyzoites was 22.89 and 53.11, respectively (Fig. 5B). The growth of acid pepsin treated Tg68 tachyzoites was evident by the average of 6.78 foci formed at day 5 post infection, while no foci were detected in the acid pepsin treated ME49 tachyzoites (Fig. 5B). These data indicate that despite forming less foci compared to bradyzoites, Tg68 tachyzoites develop resistance to acid pepsin, a functional trait of bradyzoites required for oral transmission, which aligns with the elevated basal expression of BFD1 and BFD1-regulated genes in tachyzoites of Tg68.

## Discussion

The ability of *T. gondii* to convert into bradyzoite-containing cysts within host tissues allows for long-term persistence, transmission to other hosts, and potential reactivation that causes severe disease in immunocompromised individuals(2, 44). An *in vitro* culture system for *T. gondii* cysts would aid in understanding the basis for recrudescence and possibly identify targets for intervention. Herein we characterized a natural isolate of *T. gondii* from a congenital human infection in a conventional type II strain described previously(24). Tg68 tachyzoites showed slower replication in HFFs, but robustly formed cysts and mature bradyzoites under conditions that induce differentiation including alkaline pH and low glucose. Transcriptional analysis showed that the key regulators of bradyzoite development, BFD1, BFD2, and several AP2 transcription factors that regulate bradyzoite development independently of the BFD1/BFD2 pathway, were transcriptionally altered in Tg68 tachyzoites. Consistent with these combined changes, a number of bradyzoite-specific genes were constitutively expressed in tachyzoites of Tg68. These findings indicate that Tg68 tachyzoites exist in a pre-bradyzoite state, making it permissive for bradyzoite development and cyst formation under stress conditions *in vitro* (Fig. 5C).

Previous studies have described a spontaneously differentiating strain called EGS, which is an atypical strain from South America, that has been used for studying differentiation and profiling compounds that act on bradyzoites(45, 46). The EGS strain belongs to Clade B that comprises several lineages with high virulence in laboratory mice and which is relatively distant from the commonly used lab strains(25). In contrast, Tg68 is a conventional type II strain that is highly similar to ME49 based on whole genome sequencing and phylogenetic analysis of SNPs. However, despite having low level of polymorphism, it behaves very differently phenotypically. Tg68 shows constitutive expression of bradyzoite regulatory factors, pepsin resistance as a tachyzoite, and is permissive for stress induced development of bradyzoites. Upon stress induction, Tg68 tachyzoites develop into fully differentiated bradyzoites at day 7 *in vitro*, as indicated by the comparably high expression of bradyzoite-specific genes to those found in late-stage *in vivo* bradyzoites(18).

It has long been established that bradyzoite differentiation is closely linked to a decrease in the replication rate, which is normally high in tachyzoites and considerably lower in bradyzoites(47–50). Bradyzoite conversion can be induced *in vitro* by a variety of stressors, including alkaline pH (8.2)(17), heat shock (43 °C)(17), nutrient starvation (pyrimidine and arginine starvation)(51, 52), and cholesterol deprivation(53), and as we show here, by culture in low glucose and elevated glutamine. A common feature of these stressors is their capacity to restrict tachyzoite replication. Moreover, the three major clonal lineages of *T. gondii* that have different growth characteristics, vary in their ability to form tissue cysts, reflecting an inverse association between tachyzoite replication and bradyzoite differentiation. Although RH tachyzoites have been reported to express bradyzoite-specific genes under stress conditions(17), type I strains typically exhibit rapid replication but are largely incapable of forming mature cysts *in vitro* or *in vivo*(54, 55). In contrast, type II/III strains proliferate relatively slowly and readily convert to bradyzoites under stress conditions (54, 55). Tg68 tachyzoites replicate at a significantly slower rate and are more permissive for bradyzoite differentiation under alkaline stress *in vitro* in comparison with ME49 tachyzoites, supporting the correlation between slower replication and differentiation.

Genome wide SNP analysis identified polymorphisms in Tg68 that could potentially affect parasite replication. We identified a loss in the initiating ATG in *TgGC* of Tg68. TgGC produces cyclic guanosine monophosphate (cGMP) and controls microneme secretion, that impacts motility, invasion, and egress(26). The potential use of an alternative downstream ATG may affect levels of *TgGC*, resulting in impaired lytic growth of the parasite, which may explain the observed slow lytic growth of Tg68. Alternatively, EF-Ts interacts with elongation factor thermos-unstable (EF-Tu) and mediates translation in the mitochondrion(27). The frameshift variant found in *EF-Ts* in Tg68 results in a truncation of the mitochondrial EF-Ts protein that lacks two conserved interacting residues with EF-Tu, which potentially led to a slow recycling of EF-Tu and deficient mitochondrial translations(27, 56). It has been shown that the defect in mitochondrial translation resulted in parasite growth defect(57), and the impact of the truncated EF-Ts in Tg68 on parasite growth will require further investigation. We did not identify an obvious genetic basis for the permissive bradyzoite differentiation of Tg68. In Tg68, *BFD1* contained a synonymous mutation, and no variants were detected in *BFD2*. Among the six AP2 factors that were associated with bradyzoite development, *AP2IV-3* had a variant in the upstream region of the gene; *AP2IX-9* was found to have two mutations in the 3’ untranslated region (UTR), one in the upstream region, and one synonymous mutation. Hence, it remains possible that sequence variants underlie the stable changes in gene expression observed in Tg68.

It has been postulated that a pre-bradyzoite is an intermediate stage during *T. gondii* tachyzoite-to-bradyzoite development(20, 48, 50). Jerome *et al*. showed that the fast-growing tachyzoites shift to slower growth, which preceded the expression of the bradyzoite-specific marker, *BAG1*(50). Consistently, Radke *et al*. showed a subpopulation of parasites with cell cycles of late-S/G2 arising transiently at the onset of bradyzoite differentiation(48). Here we showed that Tg68 tachyzoites shared striking similarities to such transitioning parasites in terms of slower parasite growth. Gene expression difference observed in Tg68 tachyzoite are also consistent with a pre-bradyzoite intermediate stage. Previous microarray studies have shown induction of bradyzoite genes in a progressive manner from tachyzoites to pre-bradyzoites to mature bradyzoites, including *ENO1*(20). Similar to this profile, we observed that Tg68 constitutively express highly levels of *BFD1*, *BFD2*, AP2 factors, and a constellation of bradyzoite specific genes including *ENO1*, *CST1*, *BAG1*, and many others. These findings demonstrate that Tg68 represents a pre-bradyzoite state that is primed to convert to bradyzoites following stress induction. As such, this strain may be useful in further analyzing the factors that lead to induction of altered gene expression during differentiation, including how stress pathways impact translation and transcription. Our findings also illustrate how subtle changes in conventional genotypes can have propound impacts phenotypic traits in parasites.

## Methods

### Parasite and cell culture

*T. gondii* tachyzoites were maintained by serial passage in HFFs in Dulbecco’s Modified Eagle’s Medium (DMEM; Thermo Fisher) containing 10% fetal bovine serum (FBS), 2 mM glutamine (Sigma), and 10 μg/mL gentamicin (Thermo Fisher) at 37°C in a 5% CO_2_ incubator. For *in vitro* bradyzoite differentiation under alkaline pH conditions, HFFs were infected with tachyzoites and incubated for 4 hr to allow for invasion. Culture medium was then switched to RPMI 1640 (Sigma) containing 1% FBS and 50 mM HEPES (Sigma), adjusted to pH 8.2 and the culture was incubated at 37°C in ambient CO_2_. For *in vitro* bradyzoite differentiation under glucose-free conditions, HFFs were infected with tachyzoites and incubated for 4 hr to allow for invasion. Culture medium was then switched to glucose-free RPMI 1640 (Sigma) containing 1% FBS, 50 mM HEPES (Sigma), and 10 mM glutamine (Sigma), adjusted to pH 7.2 and the culture was incubated at 37°C in 5% CO_2_. Parasite cultures and host cell lines were shown to be negative for mycoplasma using an e-Myco plus kit (Intron Biotechnology).

### gDNA isolation and whole genome sequencing (WGS)

Infected HFFs were scraped and syringe-lysed using 25 g and 27 g needles to release the parasites. The parasites were harvested by passage through a 3-μm filter and centrifugation at 400 x g for 10 min. gDNA was isolated using DNeasy Blood & Tissue Kit (QIAGEN). WGS was conducted using Illumina NovaSeq 6000 s4 system with 2×150 bp paired-end technique.

### SNP identification and annotation

Sequencing reads were trimmed for size and quality using the error rate limit setting of 0.02, maximum number of ambiguities setting of 2, and minimum read length setting of 65 nucleotides. The trimmed reads were aligned against the ME49 reference genome (ToxoDB version 59; https://toxodb.org), and single nucleotide variants and short indels were identified using CLC Genomics Workbench 21.0 with minimum coverage of 10, minimum read count of 2, and minimum variant frequency of 90%. The variants identified were annotated and analyzed using SnpEff(58).

### Phylogenetic Network analysis

Genome-wide SNPs of 62 indicated *T. gondii* strains(25) together with Tg68 were used to construct phylogenetic tree with SplitsTree 4(59) using a neighbor-net method and 1,000 replicates of bootstrapping.

### Antibodies

Anti-SAG1 mouse mAb DG52 and anti-SAG1 rabbit antibodies were kindly provided by Dr. John Boothroyd (Stanford University, Stanford, CA, USA). Anti-BAG1 mouse mAb 8.25.8 was kindly provided by Dr. Louis Weiss (Albert Einstein College of Medicine, Bronx, NY, USA). Fluorescein-conjugated Dolichos Biflorus Agglutinin (DBA) was obtained from Vector Laboratories. Anti-HA mouse mAb 16B12 was obtained from BioLegend.

### Immunofluorescence (IF) assay and microscopy

Samples were fixed with 4% formaldehyde in PBS for 15 min and blocked/permeabilized with PBS containing 5% BSA and 0.1% Triton X-100 for 30 min. For vacuolar size assay and bradyzoite differentiation assay of dual reporter strain, cells grown in black 96 well plates with flat clear bottom (Corning) were stained with Hoechst 33342 (Life Technologies) for 20 min and were imaged with a Cytation 3 Imager (BioTek). The diameter of SAG1-positive vacuoles was determined using CellProfiler 4.2.4(60). Fluorescence intensities of SAG1-EGFP and BAG1-mCherry were determined using CellProfiler 4.2.4. For bradyzoite development assay of wild type parasites and cyst purification assay, cells grown on glass coverslips in 24 well plates were incubated with indicated primary antibodies for 2 hr, followed by rinsing and incubation with Alexa Fluor-conjugated secondary antibodies for 1 hr. Cells were mounted in ProLong Gold antifade mountant (Invitrogen) and sealed onto glass slides. Cells were visualized and images were captured with a Zeiss Axioskop2 MOT Plus wide-field fluorescence microscope (Carl Zeiss, Inc.) using a ×63 oil Plan-Apochromat lens (N.A. 1.4). Images were acquired using Axiovision LE64 software (Carl Zeiss, Inc.). For focus forming assay, cells grown in black 24 well plates with flat glass bottom (Cellvis) were visualized based on green fluorescence, and foci were counted with a Zeiss Observer Z1 inverted microscope (Zeiss) using a Colibri 7 LED light source (Zeiss), ORCA-ER digital camera (Hamamatsu Photonics), Plan-Neofluar 10× (NA 0.3) objective (Zeiss), and ZEN Blue image acquisition software (v2.5).

### Construction of transgenic parasites

#### Tg68 SAG1:EGFP,BAG1:mCherry,DHFR

Fragments encoding EGFP driven by the *SAG1* promoter, mCherry driven by *BAG1* promoter, *DHFR* drug selectable marker, and pNJ-26 vector were assembled using NEBuilder HiFi DNA Assembly Master Mix (NEB), to generate the plasmid pSAG1:EGFP-DHFR-BAG1:mCherry. Tg68 tachyzoites were electroporated with 50 μg of the plasmid and subjected to pyrimethamine selection (3 μM). Stable lines were cloned by limiting dilution and confirmed for transgene expression by IFA. Primers used for constructing the strains are listed in **Table S2**.

#### BFD1-3xHA expressing strains

A Cas9-expression plasmid encoding a gRNA targeting the downstream of *BFD1* coding sequence was constructed. Parasites were transfected with 10 μg of the gRNA-Cas9-expression plasmid along with 2 μg of linearized repair template encoding 3xHA-BFD1 3’UTR and DHFR expression cassette flanked by 500 bp of homology adjacent to the gRNA target site. Transfectants were selected in pyrimethamine (3 μM) and stable lines were cloned by limiting dilution. Clones were confirmed for HA expression in *in vitro* induced bradyzoites by IFA staining for HA. Primers used for constructing the strains are listed in **Table S2**.

#### *In vitro* cyst purification

Cysts were induced under alkaline stress *in vitro* as described above and were then released from host cells by scraping and passage through a 23 g needle. Lysates were incubated with 5 μL of anti-SAG1 mAb DG52 and 50 μL of Dynabead protein G magnetic beads (Invitrogen) at 4 ℃ for 2 hr, and the unbound fraction was separated from the bound fraction using a magnet. Ten microliters of biotinylated DBA (Vector Laboratories) was added into the unbound fraction and incubated with Pierce streptavidin magnetic beads (Thermo Scientific) at 4 ℃ for 2 hr. Beads and absorbed cysts were washed three times with phosphate-buffered saline (PBS) and collected using a magnet.

### RNAseq and analysis

Total RNA was isolated using RNeasy Mini Kit (QIAGEN). Three replicates were prepared for each time point and RNA quality was assessed using an Agilent Bioanalyzer. Libraries were generated using the Clontech SMARTer low input RNA kit (TAKARA) and sequenced on the Illumina NovaSeq 6000 s4 system. Sequencing data was analyzed using Partek Flow software. Raw reads were trimmed based on minimum Phred score of 20 and minimum read length of 25 nt. Trimmed reads were aligned to the ME49 genome (ToxoDB version 59; https://toxodb.org) using STAR v2.7.8a with default settings. Differential expression analysis was conducted using DEseq2, and genes considered to be differentially expressed if the false discovery rate (FDR)-adjusted *P* value was ≤ 0.05 and absolute fold change ≥ 2.

The set of BFD1-regulated genes used for comparison in BFD1 overexpression parasites and Tg68 tachyzoites (Figure 5A) was curated from the following data: 1) BFD1 targets identified by CUT&RUN profiling from Waldman *et al*., 2020(38); 2) genes significantly affected by alkaline stress with adjusted *P* values of ≤ 0.001 in alkaline stressed versus unstressed conditions transcriptional data from Waldman *et al*., 2020(38); 3) genes significantly regulated by BFD1 overexpression with adjusted *P* values of ≤ 0.05 in Δ*BFD1/DD-BFD1-Ty* parasites+Shield-1 versus vehicle transcriptional data from Waldman *et al*., 2020(38). BFD1-regulated genes are listed in DataSet 2.

### RT-qPCR

Total RNA from parasites was isolated using RNeasy Mini Kit (QIAGEN). First-strand cDNA was synthesized using ProtoScript II first strand cDNA synthesis kit (NEB), and quantitative real-time PCR was performed on QuantStudio 3 real-time PCR system (Applied Biosystems) using PowerUp SYBR green master mix (Applied Biosystems). Relative quantitation was determined by ΔΔCt with data normalized to *GCN5B*, which is consistently expressed regardless of alkaline treatment(39). Fold changes compared to ME49 tachyzoites were derived from three independent experiments. Primers used for RT-qPCR are listed in **Table S2**.

### Acid pepsin digestion

Cysts and parasites were treated with 0.1 mg/mL pepsin (Sigma), 170 mM NaCl, 60 mM HCl at 37 ℃ for 10 min, and digestion was neutralized with 94 mM Na_2_CO_3_. Acid pepsin treated parasites were quantified using a hemocytometer and were then added on HFF monolayers. The number of foci were counted at 5 days post infection as described above.

### Statistics

Unless otherwise specified, data from three independent experiments were combined and plotted as means ± stand error of means using GraphPad Prism version 9. Statistical analyses were done using IBM SPSS Statistics version 26.0(61). Parametric statistical tests were used when *P* value of Levene’s test for equality of variance was more than 0.05. A Student’s t-test was used to determine statistical difference between two samples, and *P* values of ≤ 0.05 were considered statistically significant. Experiment-specific statistical information is provided in figure legends.

## Author Contributions

J.X.: Conceptualization, Methodology, Investigation, Data Curation, Formal analysis, Writing – Original Draft, Writing – Review & Editing, Visualization, Funding acquisition. Y.F. and W.H.: Methodology, Investigation, Writing – Review & Editing. L.D.S.: Conceptualization, Writing – Original Draft, Writing – Review & Editing, Supervision, Project administration, Funding acquisition,

## Declarations

The authors have no conflicts to declare.

## Acknowledgements

The authors thank the Genome Technology Access Center at the McDonnell Genome Institute at Washington University for technical help with genomic and transcriptional analysis. We thank Wandy Beatty, Molecular Microbiology Imaging Facility at Washington University for technical assistance with microscopy. We thank Michael White and Sebastian Lourido for helpful advice. Partially supported by a grant from the NIH to L.D.S. (AI162749) and the Stephen I. Morse Fellowship to J.X.

## Supplementary files

**Figure S1.**
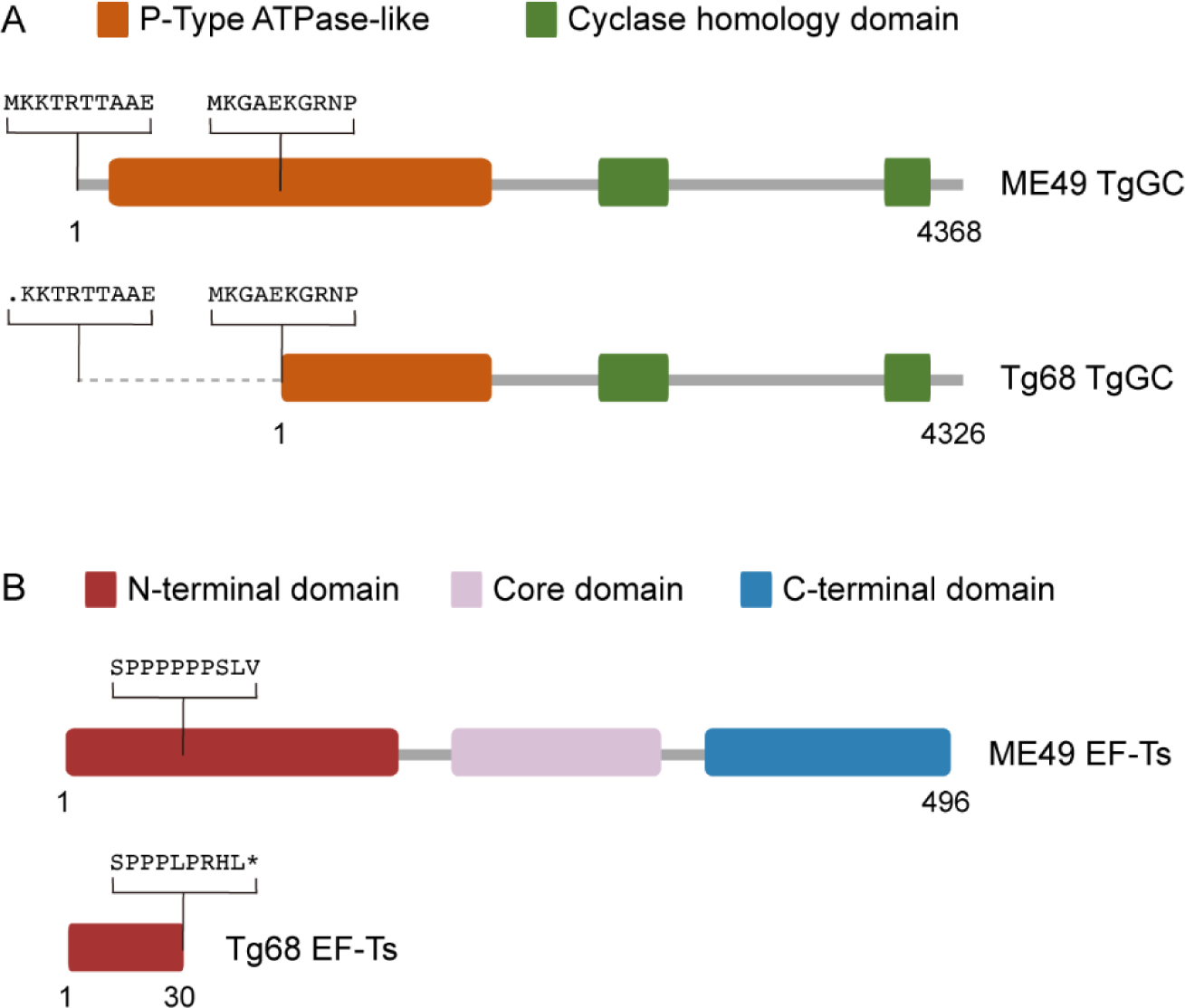
Schematic showing the variants in (A) *TgGC* and (B) *EF-Ts* in Tg68, and their potential impact on the proteins.

**Figure S2.**
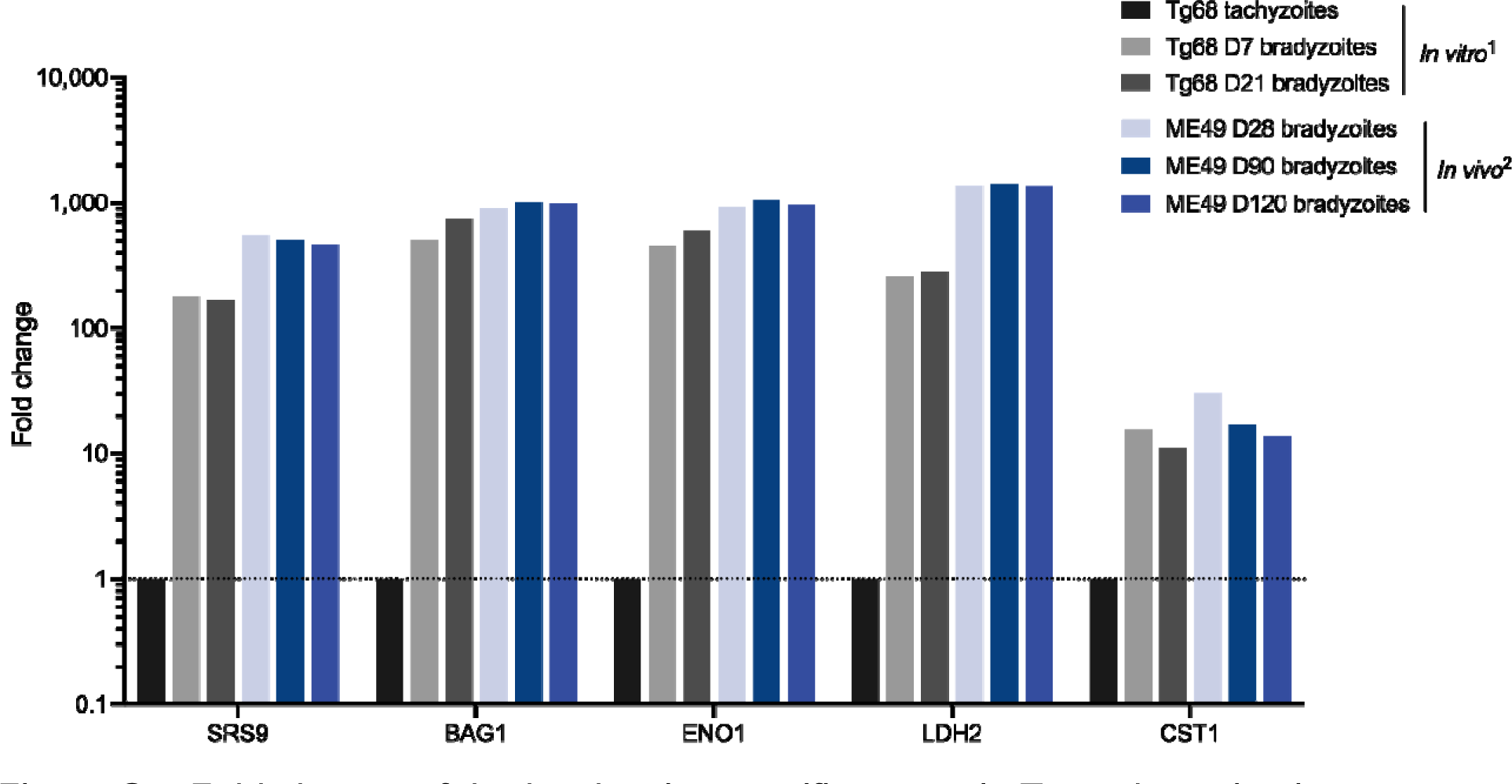
Fold change of the bradyzoite-specific genes in Tg68 day 7 *in vitro* bradyzoites, Tg68 day 21 *in vitro* bradyzoites, ME49 day 28 *in vivo* bradyzoites, ME49 day 90 *in vivo* bradyzoites, and ME49 day 120 *in vivo* bradyzoites, compared to Tg68 *in vitro* tachyzoites. ^1^Data was from this study. ^2^Data was obtained from Garfoot *et al*., 2019(18).

**Table S1** Potential functional variants identified by genome wide SNP analysis.

**Table S2** Primers used in this study.

**DataSet 1** Differential expression analysis of Tg68 tachyzoites and bradyzoites.

**DataSet 2** Gene set of BFD1-regulated genes.

